# Large *Anoura* (Chiroptera:Glossophaginae) taxonomy, taxonomic status of *Anoura carishina*, and implications for the distribution of *Anoura latidens* in Colombia

**DOI:** 10.1101/462051

**Authors:** Camilo A. Calderón-Acevedo, Miguel E. Rodríguez-Posada, Nathan Muchhala

## Abstract

The *Anoura geoffroyi* species complex is composed of 3 large species: *A. geoffroyi*, *A. peruana*, and *A. carishina.* Several inconsistencies arise from the description of *A. carishina,* and given the lack of a comparison with the dentition and external characters of *A. latidens*, here we compare the taxonomic characters of these species. To understand the position of *A. carishina* in the morphospace occupied by large *Anoura*, we conducted a Principal Component Analysis on 12 craniodental and 11 external variables. We complement our results with further analysis of traits thought to be diagnostic for these species, including 1) an elliptical Fourier transformation analysis of the shape of the third upper premolar (P4), 2) a comparison of the area of the second (P3) and third (P4) upper premolars, and 3) a comparison of maxillary toothrow angles. We find that *A. carishina* is morphologically indistinguishable from *A. latidens*, and that there is broad overlap in morphology between *A. latidens* and *A. geoffroyi*. However several characters found in *A. latidens* are lacking in *A. geoffroyi*, including a triangular shape to the P4 caused by a medial-internal cusp enclosed by the base of the tooth, a lack of development of the anterobasal cusp in the P3, a smaller braincase, and a shorter rostrum. We reassess the distribution of *Anoura latidens* in Colombia, adding new records and correcting previously-published records that were misidentified. Overall, our results suggest that a stable taxonomy for the group should consider *A. carishina* as a junior synonym of *A. latidens,* and that, although *A. latidens* is distinguishable from *A. geoffroyi*, further genetic and taxonomic work is needed in to clarify species limits within the *A. geoffroyi* species complex.

El complejo de especies *Anoura geoffroyi* se compone de 3 especies, *A. geoffroyi*, *A. peruana,* y *A. carishina.* La descripción de *Anoura carishina* posee varias inconsistencias, y dado que no se realizó una comparación con *A. latidens,* realizamos una comparación de los caracteres taxonómicos de ambas especies. Para entender la posición de *A. carishina* en el morfoespacio ocupado por los *Anoura* grandes realizamos un Análisis de Componentes Principales usando 12 variables cráneo-dentales y 11 variables externas. Complementamos nuestros resultados con 1) un análisis de transformación elíptica de Fourier de la forma del tercer premolar superior (P4) 2) una comparación del área del segundo (P3) y tercer (P4) premolares superiores y 3) una comparación de los ángulos de las hileras dentales maxilares. Encontramos que *A. carishina* es morfológicamente indistinguible de *A. latidens* y que existe una amplio superposición en la morfología de *A. latidens* y *A. geoffroyi*. Sin embargo, la forma del P4, una cúspide anterobasal no desarrollada en el P3, y caracteres relacionados con una caja craneana menos inflada y un rostro corto son útiles en distinguir *A. latidens* de *A. geoffroyi.* Reevaluamos la distribución de *Anoura latidens* en Colombia, al agregar nuevos registros y corrigiendo registros previamente publicados que se encontraban mal identificados. En general, nuestros resultados sugieren que una taxonomía estable para el grupo debería considerar a *A. carishina* como un sinónimo junior de *A. latidens,* que *A. latidens* es distinguible de *A. geoffroyi* utilizando medidas cráneo-dentales y resalta la necesidad de estudios genéticos para esclarecer las relaciones filogenéticas entre *A. latidens* y el complejo de especies *A. geoffroyi.*

---

*Anoura* is one of the most species-rich genera in the subfamily Glossophaginae. It is currently comprised of 10 species, although not all are widely accepted species (Handley 1984, Mantilla-Meluk & Baker 2006, Griffiths & Gardner 2007 [2008], Jarrín-V & Kunz 2008, Mantilla-Meluk & Baker 2010, Pacheco et al. 2018). The genus is subdivided into two groups based on dental morphology and size (Allen 1898, Griffiths & Gardner 2007 [2008]), with five small species (*A. caudifer*, *A. aequatoris*, *A. cadenai*, *A. fistulata and A. luismanueli*) and five large species (*A. carishina*, *A. cultrata*, *A. geoffroyi*, *A. peruana* and *A. latidens*). Mantilla-Meluk and Baker (2010) designated three of these large species (along with their subspecies) as the *A. geoffroyi* species complex, including *A. carishina*, *A. geoffroyi geoffroyi*, *A. geoffroyi lasiopyga* and *A. peruana,* and also elevated *A. peruana* to a separate species rather than a subspecies of *A. geoffroyi.*

*Anoura carishina* Mantilla-Meluk and Baker 2010 is only known to date from the 5 specimens of the type series deposited at the Mammal Collection Alberto Cadena García at Instituto de Ciencias Naturales (Universidad Nacional, Bogotá, Colombia). Its distribution is limited to 3 localities in the western slopes of the southern Colombian Andes and the Sierra Nevada de Santa Marta, a mountain system isolated from the Andes in the north of Colombia. The type ICN-14530 and paratype ICN-14531 are from Taminango, Nariño department (1.67°, -77.32°). The two other localities are San Pedro de La Sierra, Sierra Nevada de Santa Marta, department of Magdalena (10.90°, -74.04°) for paratypes ICN-5224, 5225 and Cali, Pance, department of Valle del Cauca (3.32°, -76.63°) for paratype ICN-5938. *Anoura carishina* was described as a large *Anoura* with the following diagnostic characters: greatest length of skull less than 24.5 mm, small canines, P4 teeth with a wide triangular base, and complete zygomatic arches (although they are broken in several of the type series collections; (Mantilla-Meluk & Baker 2010)). However, in the description it was only explicitly compared to the subspecies of *Anoura geoffroyi* (*A. g. geoffroyi*, *A. g. lasiopyga*) and *A. peruana* - it was not compared to *A. latidens*, a species to which it bears resemblance in dental morphology, size, and coloration.

*Anoura latidens* Handley 1984 is described as a large species of *Anoura,* distinguishable from *A. geoffroyi* by a relatively short rostrum, an inflated braincase, nearly parallel maxillary toothrows, and smaller and more robust premolars which have a quadrangular appearance when viewed from above. More specifically, Handley (1984) states that the third upper premolar (P4) has a medial-internal cusp enclosed in the triangular base of the tooth (rather than an abruptly protruding cusp as in *A. geoffroyi*) and that the second upper premolar (P3) possesses a reduced anterobasal cusp. The holotype is from Pico Ávila, Caracas, Venezuela, and the species has been reported for at least 14 localities in Venezuela (Handley 1976, 1984, Linares 1986, 1998), where it occupies a variety of ecosystems with an altitudinal range from 50 to 2600 meters above sea level. Outside of Venezuela *A. latidens* has only been registered in a handful of localities in Colombia, Guyana, and Peru (Handley 1984, Linares 1998, Solari et al. 1999, Lim & Engstrom 2001), suggesting a wide yet discontinuous distribution.

In Colombia, *Anoura latidens* is distributed in the Andean region (eastern, central, and western mountain ranges) and the inter-Andean valleys (Alberico et al. 2000, Solari et al. 2013). The first record for the country was mentioned in the species description (Handley 1984) as collected by Nicéforo María in 1923 in San Juan de Rioseco, department of Cundinamarca, on the western slope of the Cordillera Oriental (eastern mountain range) above the inter-Andean valley of the Magdalena river at a height of 1000 meters above sea level. Later Muñoz (2001) attributed the first record to Wilson & Reeder (1993) and added a new locality in the Cordillera Oriental (eastern mountain range) in the municipality of Gramalote, Norte de Santander department, however they did not give a catalog number for this collection supposedly located in the Museo de Ciencias Naturales de La Salle. Two other localities are reported by Rivas-Pava et al. (2007) based on three specimens deposited at Museo de Historia Natural de la Universidad del Cauca (MHNUC) from the municipalities of Acevedo (Huila department) and Argelia (Cauca department). The most recent recorded locality is Reserva Forestal Bosque de Yotoco (Valle del Cauca department) in the southwestern Andes, with one specimen deposited in the Instituto de Ciencias Naturales (ICN) mammal collection (Mora-Beltrán & López-Arévalo 2018). With only 5 localities, the knowledge of *A. latidens* in Colombia is scarce, which impacts the understanding of its conservation threats.

In this study we use morphometric approaches to reevaluate the taxonomy of the *A. geoffroyi* species complex. We focus particularly on the extent to which *A. carishina* and *A. latidens* are distinguishable from each other and other species in the complex. We also examine all known Colombian records of *A. latidens* to evaluate its distribution within the country.

## MATERIALS AND METHODS

We measured 260 individuals from the *A. geoffroyi* species complex, including 5 *A. carishina,* 48 *A. peruana,* 59 *A. latidens*, and 148 *A. geoffroyi* (106 *A. g. geoffroyi* and 42 *A. g. lasiopyga*) (See Supplementary Data SD1 for specimens reviewed and measured). We measured 12 cranial and 11 postcranial variables to the nearest 0.01 mm. Craniodental characters included: greatest length of skull (GLS, distance from the most posterior point of the skull to the most anterior point of the premaxilla not including incisors), condylobasal length (CBL, distance from the most posterior point of the condyles to the most anterior point of the premaxilla not including incisors), postorbital breadth (PB, minimum interorbital distance measured across the frontals), braincase breadth (BCB, greatest breadth of the braincase, not including the mastoid and paraoccipital processes), height of braincase (HBC, distance from the ventral border of the foramen magnum to the parietal), mastoid breadth (MB, greatest width at the mastoid processes), maxillary tooth-row length (MTRL, distance from the most posterior point of the third upper premolar to the most anterior point of the upper canine), palatal length (PL), breadth across third upper molars (M3-M3), breadth across upper canines (C-C), mandibular length (MANL, distance from the condyles to the anterior face of the mandible) and mandibular tooth-row length (MANTRL, distance from canine to the third mandibular molar). Postcranial measurements included: forearm (FA, measured from the olecranon to the articulation of the wrist), length of 3^rd^ (D3MC), 4^th^ (D4MC) and 5^th^ (D5MC) metacarpals, length of the 1^st^ and 2^nd^ phalanxes of 3^rd^ (D3P1, D3P2), 4th (D4P1, D4P2) and 5^th^ (D5P1, D5P2) digit, and length of the tibia (Tibia). Measurements were selected based on their frequent use in bat taxonomy (Handley 1960, Nagorsen & Tamsitt 1981, Handley 1984, Velazco 2005, Mantilla-Meluk & Baker 2006, Velazco & Patterson 2008, Mantilla-Meluk & Baker 2010, Velazco & Simmons 2011). Note that our measurement of the greatest length of the skull differs from that in the description of *Anoura carishina* (Mantilla-Meluk & Baker 2010) in that we measure from the posterior-most point of the occipital to the anterior-most point in the premaxilla (excluding incisors), the same measurement used in all other *Anoura* descriptions (Handley 1960, 1984, Molinari 1994, Muchhala et al. 2005), while in its description *A. carishina* and the specimens to which it was compared were measured from the posterior-most point of the occipital to the anterior-most point of the nasal bones. To analyze the morphospace of *Anoura* and explore the morphometric variation of our traits, we performed a Principal Component Analysis (PCA) for 2 data sets. One dataset (*n* = 202) includes only the 12 craniodental measurements; the second dataset (*n* = 125) includes all 23 craniodental and postcranial measurements. Both datasets include representatives of all species of *Anoura*.

To test the reliability of dental characters distinguishing *A. latidens* and *A. carishina* from *A. geoffroyi,* we traced the contour of the premolars from digital photographs of the ventral view of the skull of 70 *A. latidens, 36 A. geoffroyi*, 7 *A. peruana* and *5 A. carishina.* We took each photograph next to a band of millimeter paper in order to standardize measurements. We selected the contour of the P3 and P4 using ImageJ (Schneider et al. 2012), and obtained the area of this contour using the “Measure” function. To quantify the shape of the P4 (irrespective of size) we transformed every contour image of the P4 to a binary image in Image J (Schneider et al. 2012) and then employed an elliptical Fourier transformation on these images. Using SHAPE v1.3 (Iwata & Ukai 2002) this contour was transformed into chain code, assigning a string of code that represents the perimeter of every image of the third upper premolar, which was then used to create a harmonic or elliptical Fourier descriptor (EFDs) series. This approach allowed us to quantify the shape using 20 harmonics, which were used as input for a PCA.

Aside from tooth morphology, another character cited by Handley (1984) as important in distinguishing *A. latidens* from *A. geoffroyi* is that the former have nearly parallel maxillary toothrows. To quantify this, we used ImageJ to overlay lines over images of the occlusal view of the maxillae for 5 *A. latidens,* 34 *A. geoffroyi*, 4 *A. peruana* and 66 *A. carishina.* Specifically, these lines connected the metastyle of the third upper molar (M3) to the most anterior point of the canines for each toothrow (See Supplementary Data SD 3, Fig. 3). We then measured the angle between these lines.

We tested for significant differences between *A. geoffroyi*, *A. latidens*, *A. peruana* and *A. carishina* in 1) craniodental measurements (including those related to rostrum length and an inflated braincase) 2) P4 and P3 size (e.g. total surface area), 3) the shape of P4 (EFD principal components) and 4) the toothrow angle using a Multivariate Analysis of Variance (MANOVA) followed by Bonferroni-corrected posthoc tests for each variable.

To assess the geographical distribution of *A. latidens* we reviewed the published records and examined the skulls of specimens labeled as *A. geoffroyi* and *A. caudifer* in the following collections: Colección de Mamíferos Alberto Cadena García at Instituto de Ciencias Naturales de la Universidad Nacional de Colombia (ICN), Instituto de Investigación en Recursos Biológicos Alexander von Humboldt (IAvH), Museo Universidad Distrital Francisco José de Caldas (MUD), Museo de Historia Natural de la Universidad del Cauca (MHNUC), Colección Teriológica Universidad de Antioquia (CTUA), National Museum of Natural History (USNM), Muséum d’Histoire Naturelle de la Ville de Genève (MHNG), American Museum of Natural History (AMNH), and Field Museum of Natural History (FMNH).

## RESULTS

### Morphological revision

The type specimen of *A. carishina* (ICN 14530) evidences the dental characters provided in the description of *A. latidens* (Handley 1984). It has broad molars and premolars with the anterobasal cusp of the second upper premolar (P3) reduced and the medial-internal cusp of the third upper premolar (P4) enclosed in a triangular base. When comparing the type of *A. latidens* to the type series of *A. carishina* we find that specimens ICN 14530,14531, 5224 and 5225 possess both characteristics, while specimen ICN 5838 possesses neither and is instead diagnosable as *A. geoffroyi* (Fig.1). In our review of the type material, we also discovered that the specimen labeled as the holotype in Figure 4 of Mantilla-Meluk and Baker (2010) is in fact ICN-5225, while the specimen labeled as ICN-5225 is actually the type (ICN-5225 is a female paratype that possesses both auditory bullae, while ICN 14530 is a male specimen lacks the right auditory bulla; see Supplementary Data SD 3, Supplementary Fig. 1).

**Fig. 1.**
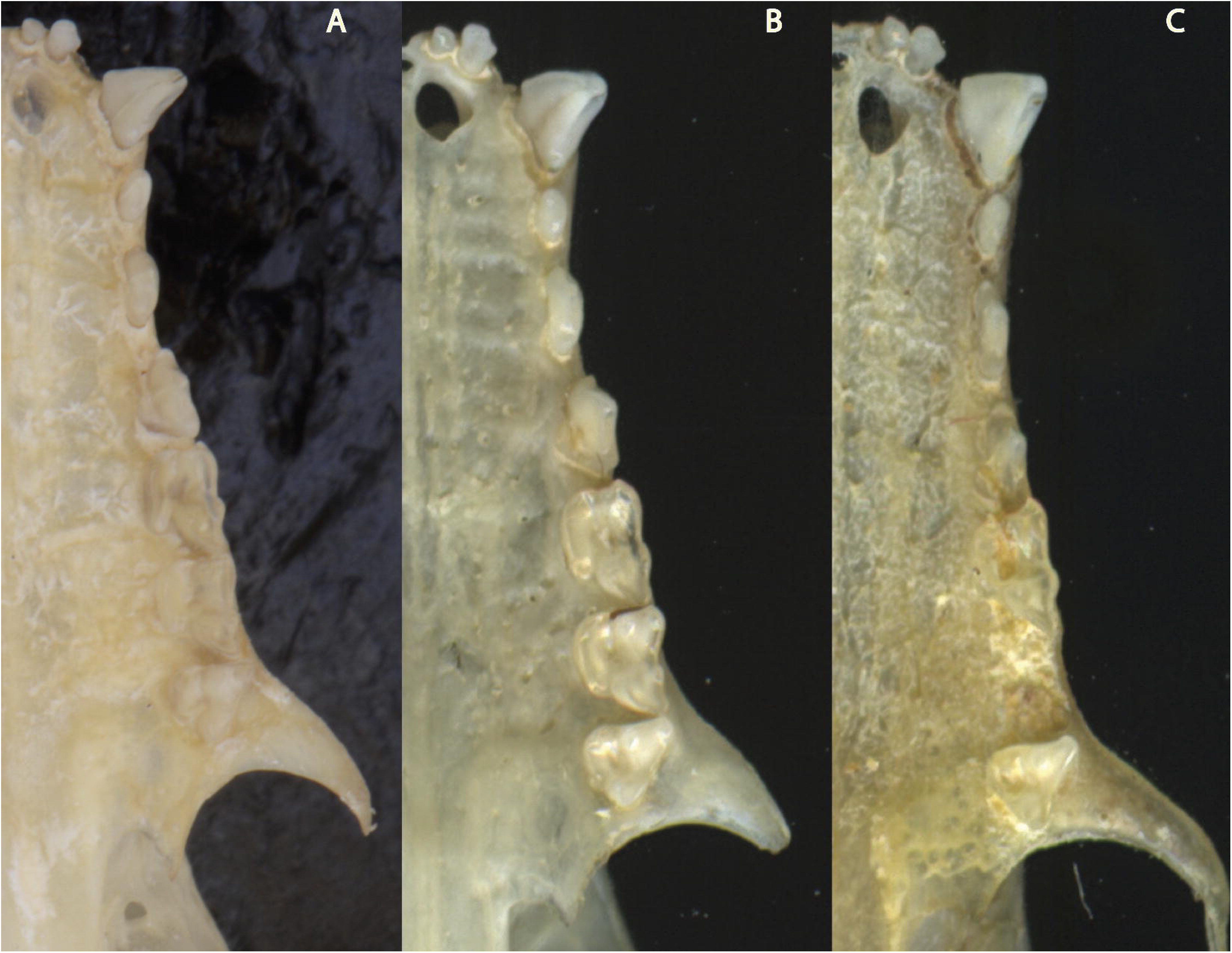
Skull morphology of A) *A. latidens* type AMNH 370119, B) *A. carishina* type ICN 14530 and C) *A. carishina* paratype ICN 5938. Note the robust molars and premolars in the first two, in contrast to the slender premolars of the *A. carishina* paratype ICN 5938.

In our review of previously-published records of *Anoura latidens* in Colombia, we find that only 2 are valid, including specimen AMNH-69187 used in the species description (Handley 1984) and ICN 22807 from Reserva Forestal Bosque de Yotoco, municipality of Yotoco, department of Valle del Cauca (Mora-Beltrán & López-Arévalo 2018). The *A. latidens* specimens reported by Rivas-Pava et al. (2007) from the municipalities of Acevedo (department of Huila; MHNUC-M0722, 0723) and Argelia (department of Cauca; MHNUC-M1552) actually correspond to individuals of *A. geoffroyi*, while there is no record of the *A. latidens* specimen reported by Muñoz (2001) in the mammal collection of Colegio San Jose de la Salle. The two putative records of *A. latidens* that we did find in this collection were both captured in Gramalote (Norte de Santander, Colombia) and are diagnosable as *Glossophaga soricina.*

On the other hand, among all of the collections we reviewed we found a total of 3 *Anoura latidens* specimens that were misidentified as other *Anoura* species. Specimens ICN 4398, ICN 11195, and MUD 587 coincide with the dental characters of *A. latidens* proposed by Handley (1984). ICN 4398 is an adult male, preserved as a skin and extracted skull. This record is located in the inter-Andean valley of the Cauca River, between the Cordillera Central and Cordillera Occidental (central and western mountain ranges). ICN 11195 is an adult male, preserved as a skin and extracted skull. It was collected in Parque Regional Natural Ucumarí, Vereda la Suiza, city of Pereira, department of Risaralda. This locality is situated in the protected area Santuario de Fauna y Flora Otún Quimbaya and resides in the western slope of the Cordillera Central (central mountain range) at an elevation of 1900 meters. MUD 587 is an adult male, preserved as a skin and extracted skull. It was collected in Vereda La Huerta, municipality of La Vega, department of Cundinamarca on the western slope of the Cordillera Oriental (eastern Andes) at an elevation of 980 meters (see Supplementary Data SD1).

### Morphometric analyses

The type series of *A. carishina* overlaps with both *A. latidens* and the *A. geoffroyi* species complex (*A. g. geoffroyi*, *A. g. lasiopyga* and *A. peruana*) in most of its measurements (Supplementary Data SD2). For the dataset with all measurements (Fig 2. A), our principal component analysis shows that less than 50% of the variation is explained by the first two principal components of the PCA (PC1 33.2%, PC2 10.7%). We recover similar results when only craniodental measurements (Fig 2. B) are taken into account (PC1 40 %, PC2 17.2%). Most of the morphospace of *A. latidens* and the *A. geoffroyi* species complex is shared in both datasets (Fig. 2, see Supplementary Data 3, supplementary Fig. 1 for the distribution of *A. g. geoffroyi*, *A. g. lasiopyga* and *A. peruana* in the morphospace).

**Fig. 2.**
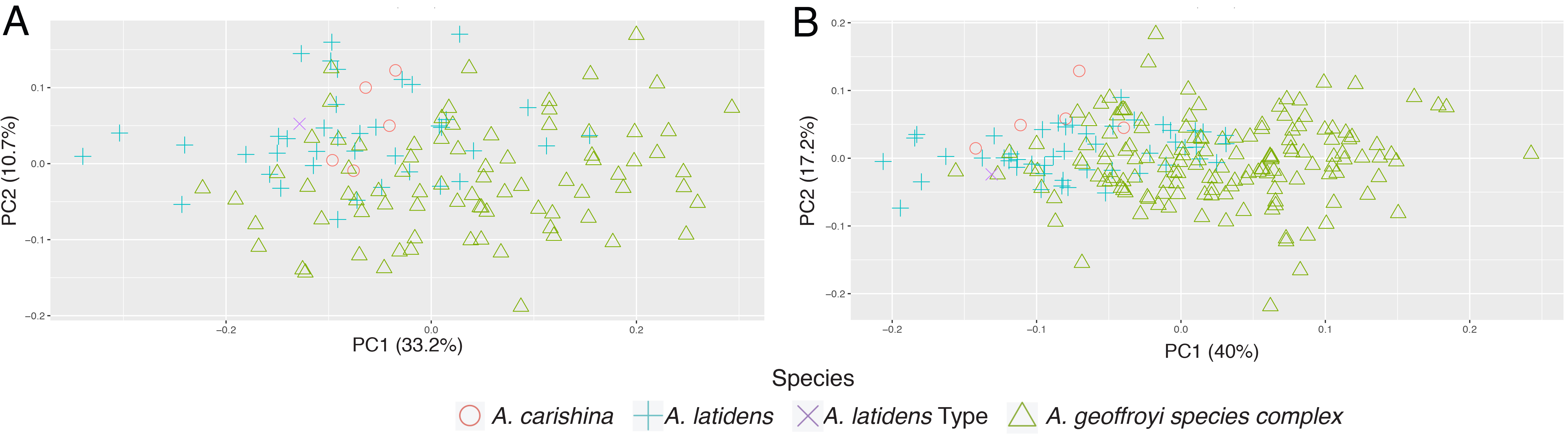
A) PCA analyses using 12 craniodental and 11 postcranial measurements of *Anoura* specimens. B) PCA analyses using only the 12 craniodental measurements of *Anoura carishina*, *A. latidens* and *A. geoffroyi* species complex specimens.

P4 shape (PCA on 20 EFDs) resulted in over 80% of the variation in the shape of the P4 (Fig. 3) being explained by the first two principal components (PC1 71.83% and PC2 13.07 %). We see that the type specimen of *Anoura carishina* (ICN 14530) is in the center of the morphospace occupied by *A. latidens*, with the position of the *A carishina* paratype diagnosable as *A. g. geoffroyi* (ICN 5938) closer to the morphospace of *A. g. geoffroyi*. Despite evidencing different morphological clusters corresponding to *A. g. geoffroyi* (with *A. peruana* immersed in its morphospace) and *A. latidens*, the morphospace of the shape of P4 does not show a clear separation between them, with some specimens of *A. g. geoffroyi, A. peruana* and *A. latidens* occupying the space between clusters (Fig. 3).

**Fig. 3.**
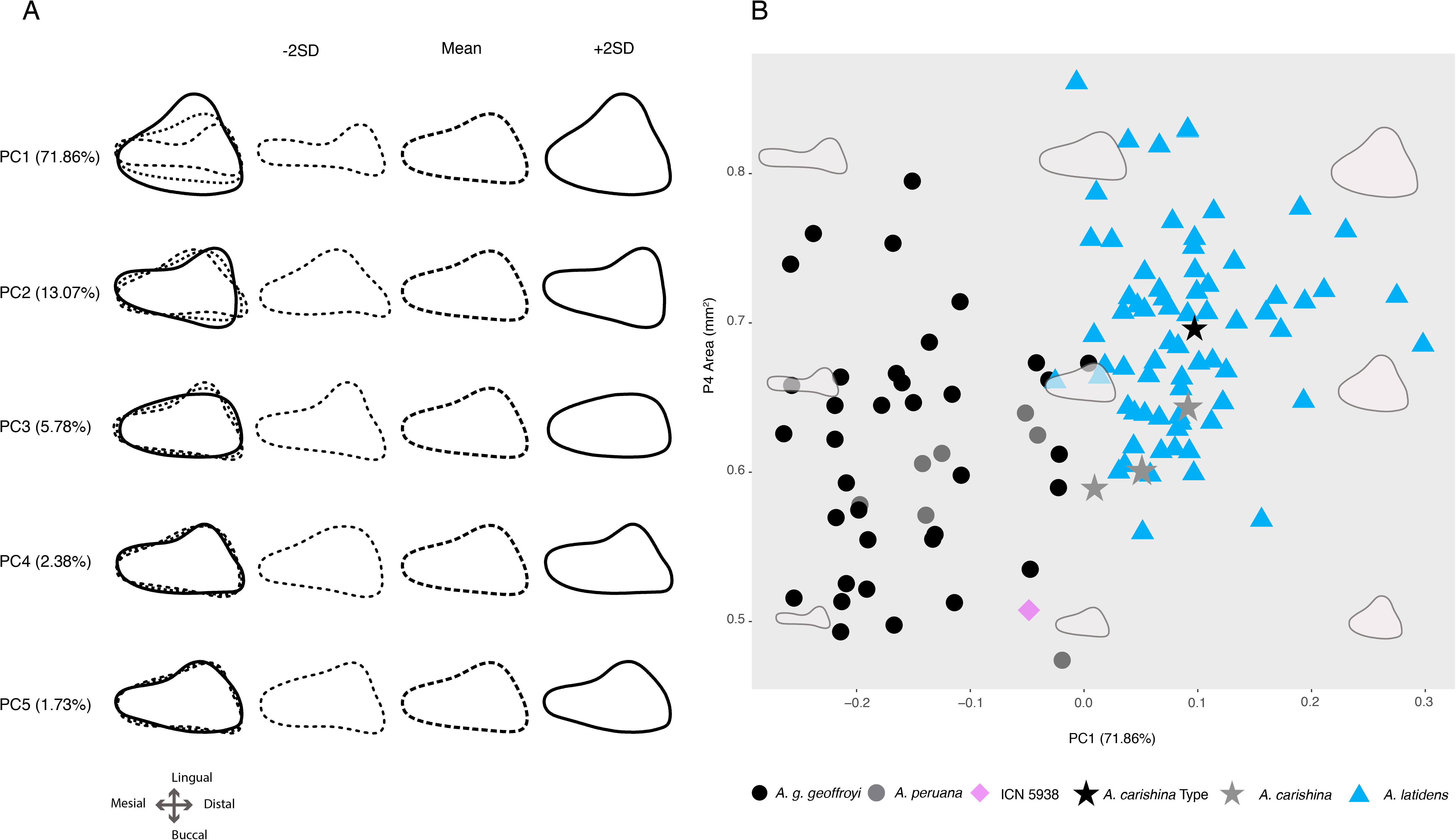
A) Mean (solid lines), -2SD (short-dashed lines), and + 2SD (long-dashed lines) contour shapes of the third premolar (P4) in our sample (with all three super-imposed to the left), showing the variation explained by each of the elliptical Fourier descriptor (EFD) Principal Components. B) Scatterplot of EFD PC1 vs. P4 area. Note that the *A. carishina* type specimen (ICN 14530) is nested well within the morphospace of *A. latidens.*

The Multivariate analysis of variance (MANOVA) of morphometric measurements showed overall significant differences for each measurement (Pillai’s Trace and Wilks’ Lamda *P*<0.001) with the exception of postorbital breadth (PB; *F_3,121_*=1.023, *P*=0.385) and forearm length (FA; *F_3,121_*=0.223, *P*=0.881) (Table 2). Bonferroni corrected *P* values show significant differences between *A. latidens* and *A. carishina* only in height of braincase (HBC; *P*=0.030), while *A. g. geoffroyi* and *A. latidens* have significant differences in the means of all except postorbital breadth (PB; *P*=1.0), height of braincase (HBC; *P*=0.166), and forearm length (FA; *P*=1.0). Of particular note are significant differences in measurements related to the overall shorter rostrum and less inflated braincase of *A. latidens*, as these features were highlighted by Handley (1984) in the description of this species. Specifically, *A. latidens* has a shorter greatest length of skull (GLS), palate length (PL), maxillary toothrow length (MTRL), braincase breadth (BCB) and mastoid breadth (MB) in comparison to *A. geoffroyi* and *A. peruana* (see Table 2, SD2). Between these latter two species, *Anoura peruana* only showed significant differences with *A. geoffroyi* in height of braincase (HBC; *P*=0.043).

**Table 1.**
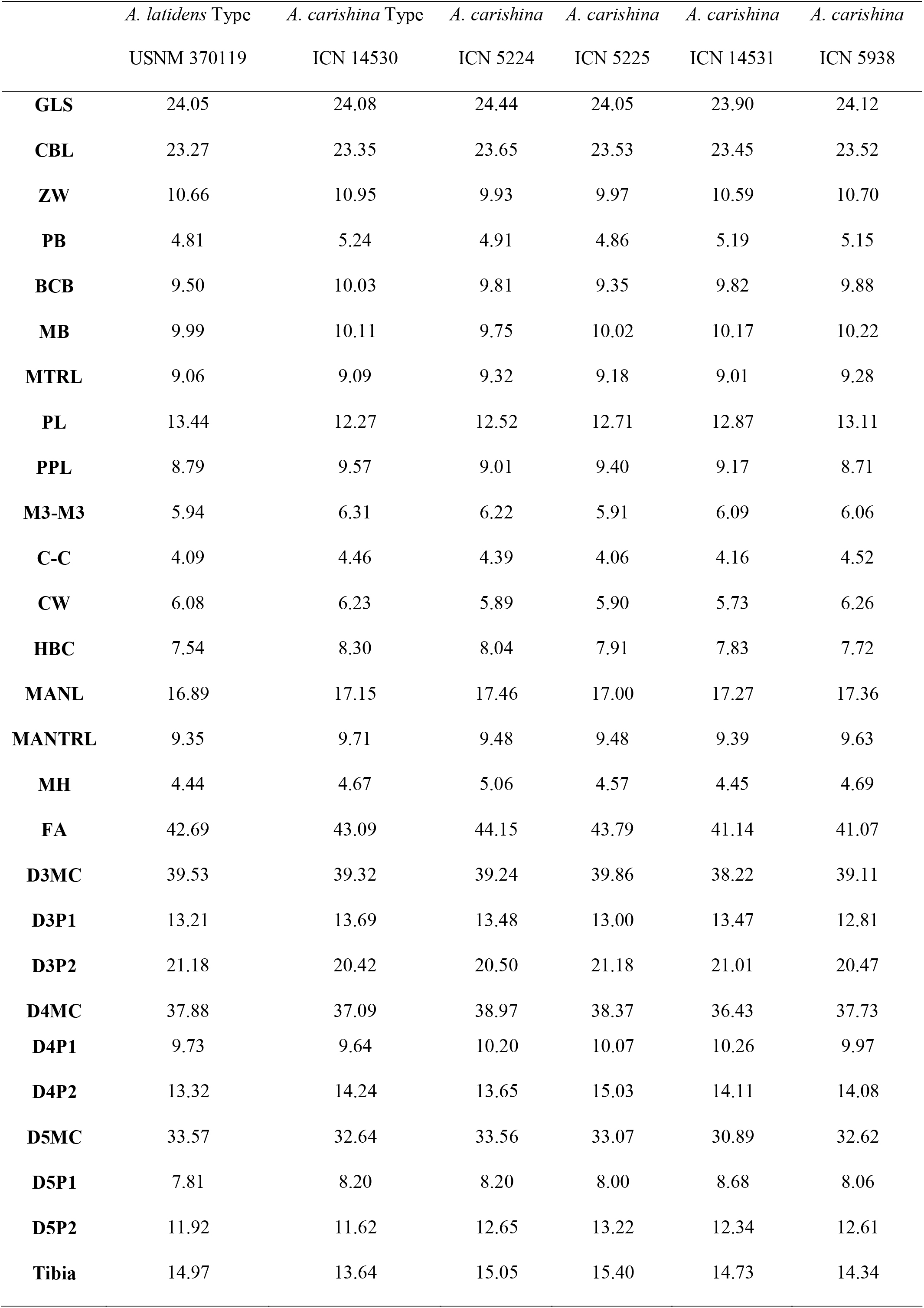
Measurements (mm) of the type specimen of *A. latidens*, and the type series of *A. carishina*, see methods for measurement abbreviations.

**Table 2.** MANOVA *F* values and *P*-values for Bonferroni-corrected posthoc tests of morphometric variables between *Anoura peruana* (*n=*5), *A. carishina* (*n*=5)*, A. geoffroyi* (*n*=75) and *A. latidens* (*n=*40), with significant *P*-values in bold. See methods for measurement abbreviations.

| Variable | MANOVA *F* | MANOVA *P* | *A. latidens* -.<br>*A. carishina* | *A. geoffroyi* -.<br>*A. carishina* | *A. peruana* -.<br>*A. carishina* | *A. geoffroyi* -<br>*A. latidens* | *A. peruana* -<br>*A. latidens* | *A. geoffroyi* -<br>*A. peruana* |
| --- | --- | --- | --- | --- | --- | --- | --- | --- |
| GLS | 33.013 | 0.000 | 1.000 | 0.001 | 0.001 | 0.000 | 0.000 | 1.000 |
| CBL | 25.771 | 0.000 | 1.000 | 0.001 | 0.006 | 0.000 | 0.001 | 1.000 |
| PB | 1.023 | 0.385 | 1.000 | 1.000 | 0.867 | 1.000 | 1.000 | 0.607 |
| BCB | 5.587 | 0.001 | 1.000 | 1.000 | 1.000 | 0.001 | 1.000 | 0.354 |
| HBC | 5.625 | 0.001 | 0.030 | 0.295 | 0.005 | 0.166 | 0.500 | 0.043 |
| MB | 9.297 | 0.000 | 1.000 | 0.047 | 1.000 | 0.000 | 1.000 | 0.255 |
| PL | 21.262 | 0.000 | 1.000 | 0.001 | 0.001 | 0.000 | 0.000 | 0.787 |
| MTRL | 9.982 | 0.000 | 1.000 | 0.087 | 0.120 | 0.000 | 0.003 | 0.415 |
| M3.M3 | 3.094 | 0.030 | 1.000 | 1.000 | 1.000 | 0.021 | 0.902 | 1.000 |
| C.C | 17.085 | 0.000 | 1.000 | 0.058 | 1.000 | 0.001 | 1.000 | 0.387 |
| MANL | 5.034 | 0.003 | 1.000 | 0.515 | 0.211 | 0.009 | 0.850 | 1.000 |
| MANTRL | 14.744 | 0.000 | 1.000 | 0.012 | 0.002 | 0.000 | 0.000 | 0.417 |
| FA | 0.223 | 0.881 | 1.000 | 1.000 | 1.000 | 1.000 | 1.000 | 1.000 |

Our MANOVA on premolar shape and toothrow angle (Table 3) showed significant differences between species in the area of P4 (*F_3,105,_=* 14.878*, P*<0.001), PC1 of P4 shape (EFDs; *F_3,105_=*103.508, P<0.001) and toothrow angles (TRA, *F_3,105_=*3.157, P=0.028). Bonferroni-corrected posthoc tests show that *A. latidens* has a larger P4 area (X□= 0.69 mm^2^) than *A. carishina* (X□= 0.61 mm^2^, *P*=0.049), *A. g. geoffroyi* (X□= 0.61 mm^2^, *P*<0.001), and *A. peruana* (X□= 0.56 mm^2^, *P*=0.002). The first principal component of the P4 shape showed significant differences between *A. g. geoffroyi* and both *A. carishina* and *A. latidens,* and between *A. peruana* and *A. latidens (P<*0.001), while *A. peruana* was not different from *A. g. geoffroyi* (*P*=0.112) or *A. carishina* (*P*=0.079). Notably, *A. carishina* is not significantly different from *A. latidens* for any of these traits except P4 area, and the four specimens of *A. carishina* diagnosable as *A. latidens* fall completely within the range of *A. latidens* variation in P4 area (Fig. 3). Toothrow angle was significantly different overall between species, however none of the Tukey nor Bonferroni corrected posthoc tests were significant between specific pairs of species (Table 3).

**Table 3.** MANOVA *F* and *P*-values for Bonferroni-corrected posthoc tests of P3 and P4 area, toothrow angles (TRA) and Principal components 1 and 2 of P4 shape between *Anoura peruana* (*n*=4), *A. carishina* (*n*=5)*, A. g. geoffroyi* (*n*=34) and *A. latidens* (*n=*66), with significant *P*-values in bold. See methods for measurement abbreviations.

| Variable | MANOVA *F* | MANOVA *P* | *A. latidens* -<br>*A. carishina* | *A. g. geoffroyi* -<br>*A. carishina* | *A. peruana* -<br>*A. carishina* | *A. g. geoffroyi* -<br>*A. latidens* | *A. peruana* -<br>*A. latidens* | *A. g. geoffroyi* -<br>*A. peruana* |
| --- | --- | --- | --- | --- | --- | --- | --- | --- |
| **P3 area** | 0.952 | 0.418 | 1.000 | 1.000 | 1.000 | 0.641 | 1.000 | 1.000 |
| **P4 area** | 14.878 | **0.000** | **0.049** | 1.000 | 1.000 | **0.000** | **0.002** | **1.000** |
| **P4 Shape PC1** | 103.508 | **0.000** | 0.678 | **0.000** | 0.079 | **0.000** | **0.000** | 0.122 |
| **P4 Shape PC2** | 0.340 | 0.797 | 1.000 | 1.000 | 1.000 | 1.000 | 1.000 | 1.000 |
| **TRA** | 3.157 | **0.028** | 1.000 | 1.000 | 1.000 | 0.066 | 0.407 | 1.000 |

## DISCUSSION

Upon revision of the type material of *Anoura carishina* and *A. latidens* we find that the type series of *A. carishina* is a mixed series of four individuals corresponding to *A. latidens* and one to *A. g. geoffroyi.* Our analyses of craniodental measurements and premolar shape of individuals of all species and subspecies in the *Anoura geoffroyi* complex (*A. geoffroyi*, *A. latidens*, *A. carishina*, and *A. peruana*) find no support for considering *Anoura carishina* as an entity morphologically distinct from *A. latidens*. Our results also clarify the characters that distinguish *A. latidens* from *A. geoffroyi*, expand the known distribution of *A. latidens* in Colombia, and raise issues regarding the conservation of this species in the country.

*Taxonomic identity of* A. carishina*—* Our different lines of evidence lead us to formally treat *Anoura carishina* as a junior synonym of *A. latidens.* First, the triangular base of the third upper premolar P4 of the type specimen of *A. carishina* (ICN 14530) is indistinguishable from *A. latidens*, as demonstrated by our analyses of tooth shape (Fig. 3), as are those of three of the paratypes. Second, we find all four of these specimens also lack a developed anterobasal cusp in the second upper premolar (P3). And finally, none of the 18 morphological measurements differ between *A. latidens* and the *A. carishina* specimens (Table 2 and 3) with the exception of the height of the brain case (HBC; *P*=0.030) and P4 area (*P*=0.049), and in both of these cases there is still extensive overlap in the range of measurements (HBC: 7.14-8.07 mm for *A. latidens* vs. 7.72-8.30 mm for *A. carishina*; P4 area: 0.56-0.86 mm^2^ for *A. latidens* vs. 0.50-0.70 mm^2^ for *A. carishina*). Given the above evidence, the holotype and three of the paratypes are diagnosable as individuals of *A. latidens*. The fourth paratype (ICN 5938) falls within the morphospace of *A. g. geoffroyi* and presents a developed anterobasal cusp in the second upper premolar, supporting its identification as *A. geoffroyi*.

*Diagnosis of* A. latidens *and* A. geoffroyi*—* Our morphometric analysis of craniodental measurements shows that *A. latidens* shares morphospace with *A. g. geoffroyi* and *A. peruana*. Of the traits mentioned by Handley (1984) to diagnose *A. latidens* from *A. geoffroyi,* we found several to still be reliable in our larger dataset in separating *A. latidens* from the *A. geoffroyi* species complex, including a more robust and more triangular third upper premolar (P4; see Fig. 3), a reduced anterobasal cusp of second upper premolar (P3), and a shorter rostrum (in terms of GLS, PL, MANL; Table 2, Supplementary Data SD2). We add to this list mastoid breadth (MB) and mandibular tooth row length (MANTRL), which are also smaller for *A. latidens* (Table 2, Supplementary Data SD2). Toothrow angle, which Handley (1984) suggested is more parallel for *A. latidens*, did not in fact show significant differences (after Tukey and Bonferroni corrected posthoc tests) between any of the species in our analyses (Table 3). Finally, although Handley (1984) suggested that *A. latidens* has a more inflated braincase, we found that its braincase (BCB, Table 2, Supplementary Data SD2) is in fact significantly less inflated than *A. geoffroyi* and *A. peruana*.

*Distribution and implications for the conservation of* Anoura latidens *in Colombia —* By combining the 2 valid previously-published records of *Anoura latidens* in Colombia (Handley 1984, Mora-Beltrán & López-Arévalo 2018) with the 7 records we found here, we report *A. latidens* in 7 localities across the country (Fig. 4, Supplementary Data SD1). With the exception of the Sierra Nevada de Santa Marta, all localities fall within highly altered ecosystems (IAvH 2004). Vereda El Hormiguero (ICN 4398) is located in a sugar cane agricultural system, even at the time of the capture of the specimen (Arata et al. 1967). San Juan de Rioseco (AMNH 69187) and Vereda La Huerta (MUD 587) are mountainous areas with a landscape composed of ranching pastures, small agricultural fields, and fragments of natural forests. Vereda La Suiza (ICN 11195) presents a heterogeneous forest cover composed of fragments of natural forests, secondary forests, and reforested areas; it is part of the Santuario de Fauna y Flora Otún Quimbaya, registered in the Colombian National System of Protected Areas (SINAP) (Estrada-Villegas et al. 2010). Reserva Forestal Bosque de Yotoco (ICN 22807) is a protected reserve in the Valle del Cauca department on the eastern slopes of the Western Cordillera. All records are located in the Andean region and the Sierra Nevada de Santa Marta between 590 and 1690 m.a.s.l. (Fig. 4, Supplementary Data SD1). In Venezuela, *A. latidens* has a similar elevational distribution, with records from 50 to 2240 meters above sea level and the majority (81%) located between 1000-1500 m.a.s.l. (Handley 1984, Linares 1986, Soriano et al. 2002).

**Fig. 4.**
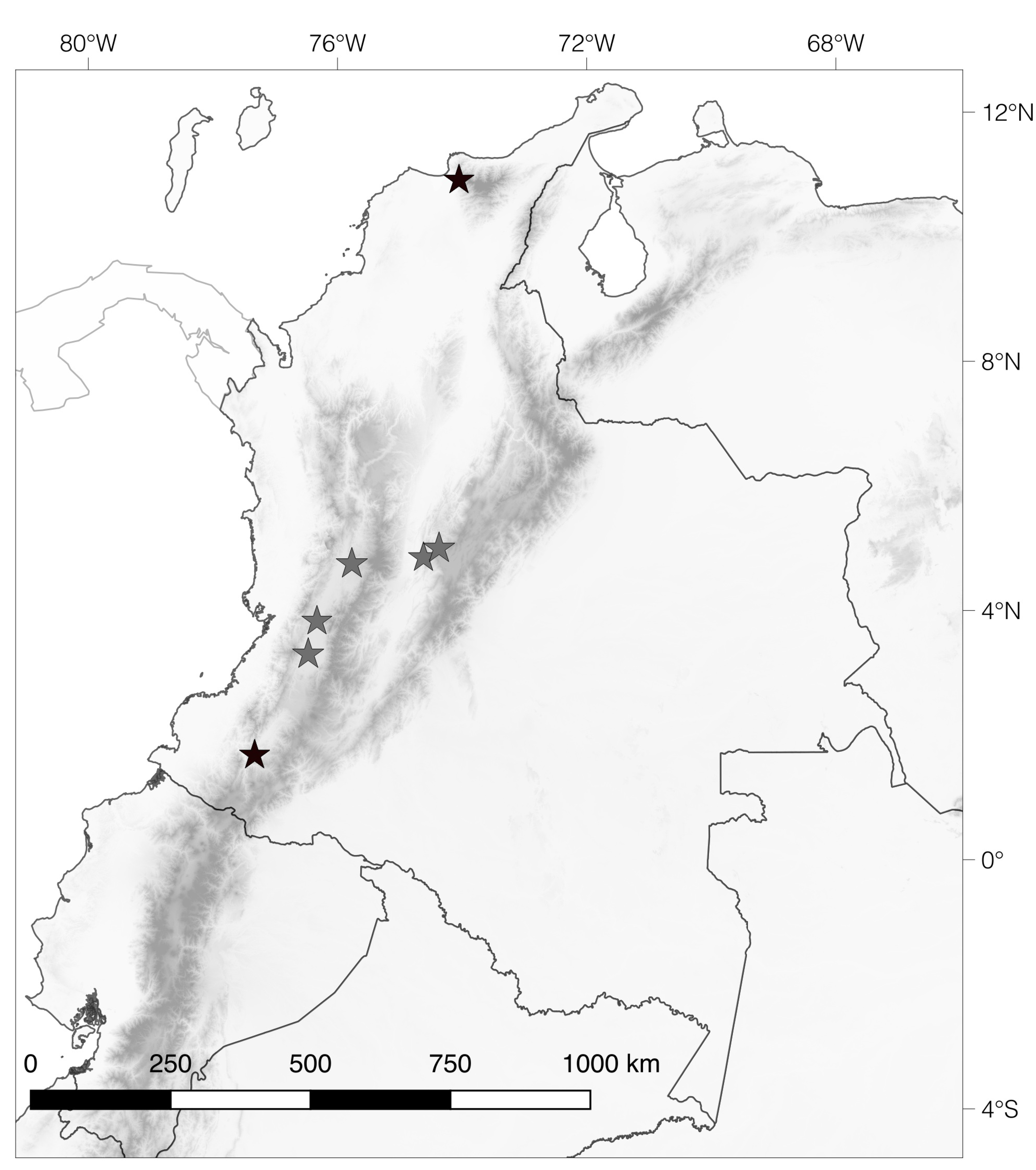
Distribution of *A. latidens* in Colombia. Black stars show specimens previously attributed to *A. carishina*, while grey stars show all other records.

Assessing the conservation status of *A. latidens* in Colombia under the conventional parameters (variation in population size, size of distribution range and habitat loss) becomes a challenge given its discontinuous distribution. It is immersed in highly transformed environments and not associated with natural vegetation cover. Local abundances are also unknown, but its limited presence in Colombian mammal collections suggests a pattern of low abundance in the Colombian Andes. Adding to this issue, given that *A. latidens* is sympatric to *A. geoffroyi*, and only craniodental features are useful for its diagnosis, it is likely that they are misidentified during fieldwork, as suggested by the fact that all new records for Colombia were previously identified as *A. geoffroyi*. In summary, *Anoura latidens* is a species with a broad distribution with unknown population numbers inhabiting highly transformed ecosystems. It is crucial to coordinate strategies with the different Bat Conservation Programs in South America to encourage research and conservation on this species, which can lead to effective conservation strategies.

In conclusion, this study provides evidence that *A. carishina* should be treated as a junior synonym of *A. latidens*, given extensive overlap in morphology, including key traits such as 1) shape of the upper third premolar (P4), 2) craniodental measurements and 3) the presence of the anterobasal cusp in the second upper premolar (P3). We found support for several characters suggested by Handley (1984) to distinguish *A. latidens* from *A. geoffroyi*, including a shorter rostrum, more robust premolars, and triangular shape to P4 (with medial-internal cusp being enclosed by the base of the tooth), while we detected no differences in toothrow angle. Finally, contrary to Handley (1984), we find that the braincase of *A. latidens* is in fact significantly less inflated than that of *A. geoffroyi*. Given the high morphological overlap between *A. geoffroyi* subspecies and *A. peruana*, we recommend further taxonomic work combined with genetic analyses to better understand the species limits of this species complex.

## ACKNOWLEDGMENTS

This work was possible thanks to the collaboration of the collection managers of multiple mammalogical collections, who provided access to specimens and their internal databases and/or provided scanned images of specimens. We want to thank H. López Arévalo (ICN), M. Ruedi (MHNG), C.A. Medina Uribe (IAVH), M. Rivas-Pava (MHNUC), A. Rodríguez-Bolaños (UD), D. Zurc (CSJ), S. Solari (CTUA), B. Patterson (FMNH), D. Lunde (USNM), and N. Simmons (AMNH). I. Jimenez provided helpful advice regarding elliptical Fourier transformation analysis.

## SUPPLEMENTARY DATA

Supplementary Data SD 1—Database of specimens examined and their geographical information including localities and geographical coordinates. Specimens revised and identified but not measured are indicated with an asterisk (^*^)

Supplementary Data SD 2—Summary measurements of *A. carishina, A. g. geoffroyi*, *A. g. lasiopyga*, *A. peruana* and *A. latidens.*

Supplementary Data SD 3— Supplementary Figure 1 Type Series of *A. carishina*, A) Type specimen ICN 14530, B) ICN 14531, C) ICN 5224, D) ICN 5225 E) ICN 5398. Supplementary Figure 2. PCA analyses discriminating between the different species/subspecies of the A. geoffroyi species complex, Top) using 12 craniodental and 11 postcranial measurements Bottom) using only the 12 craniodental measurements. Supplementary Figure 3. Depiction of toothrow angle measurement.

## LITERATURE CITED

Alberico, M., A. Cadena, J. Hernández-camacho, and Y. Muñoz-saba. 2000. Mamíferos (Synapsida: Theria) de Colombia. Biota colombiana 1:43–75.

Allen, H. 1898. On the Glossophaginae. Transactions of the American Philosophical Society 19:237–266.

Arata, A. A., J. B. Vaughn, and M. E. Thomas. 1967. Food habits of certain Colombian bats. Journal of Mammalogy 48:653–655.

Estrada-villegas, S., J. Pérez-torres, and P. R. Stevenson. 2010. Ensamblaje de murciélagos en un bosque subandino colombiano y análisis sobre la dieta de algunas especies. Mastozoología neotropical 17:31–41.

Griffiths, T. and A. Gardner. 2007 [2008]. Subfamily Glossophaginae Bonaparte, 1845. Pp. 224–244 in Mammals of South America, Vol 1 Marsupials, Xenarthrans, Shrews, and Bats (Gardner, A ed.), The University of Chicago Press, Chicago.

Handley, C. O., JR 1960. Description of new bats from Panama. Smithsonian Institution.

Handley, C. O., JR 1976. Mammals of the Smithsonian Venezuelan project. Brigham Young University Science Bulletin-Biological Series 20:1–91.

Handley, C. O., JR 1984. New species of mammals from northern South America: a long-tongued bat, genus Anoura Gray. Proceedings of the Biological Society of Washington 97:513–521.

Iwata, H. and Y. Ukai. 2002. SHAPE: a computer program package for quantitative evaluation of biological shapes based on elliptic Fourier descriptors. Journal of Heredity 93:384–385.

Jarrín-v, P. and T. H. Kunz. 2008. Taxonomic history of the genus Anoura (Chiroptera: Phyllostomidae) with insights into the challenges of morphological species delimitation. Acta Chiropterologica 10:257–269.

Lim, B. K.and M. D. Engstrom. 2001. Species diversity of bats (Mammalia: Chiroptera) in Iwokrama Forest, Guyana, and the Guianan subregion: implications for conservation. Biodiversity and Conservation 10:613–657.

Linares, O. 1986. Murciélagos de Venezuela. Cuadernos Lagoven, Caracas.

Linares, O. 1998. Mamíferos de Venezuela. Sociedad Conservacionista Audubon de Venezuela, Caracas, Venezuela.

Mantilla-Meluk, H. and R. J. Baker. 2006. Systematics of small Anoura (Chiroptera: Phyllostomidae) from Colombia, with description of a new species. Ocassional Papers, Museum of Texas Tech University 261:1–18.

Mantilla-Meluk, H. and R. J. Baker. 2010. New species of Anoura (Chiroptera: Phyllostomidae) from Colombia, with systematic remarks and notes on the distribution of the A. geoffroyi complex. Occasional Papers, Museum of Texas Tech University 292:1–19.

Molinari, J. 1994. A new species of Anoura (Mammalia Chiroptera Phyllostomidae) from the Andes of northern South America. Tropical Zoology 7:73–86.

Mora-beltrán, C. and H. F. LóPez-Arévalo. 2018. Interactions between bats and oral resources in a premontane forest, Valle del Cauca, Colombia. Therya 9:129–136.

Muchhala, N., P. V. Mena, and L. V. Albuja. 2005. A new species of Anoura (Chiroptera: Phyllostomidae) from the Ecuadorian Andes. Journal of Mammalogy 86:457–461.

Muñoz, J. 2001. Los murciélagos de Colombia. Sistemática, distribución, descripción, historia natural y ecología. Editorial Universidad de Antioquia, Medellín, Antioquia, Colombia.

Nagorsen, D. and J. Tamsitt. 1981. Systematics of Anoura cultrata, A. brevirostrum, and A. werckleae. Journal of Mammalogy 62:82–100.

Pacheco, V., P. SáNchez-vendizú, and S. Solari. 2018. A New Species of Anoura Gray, 1838 (Chiroptera: Phyllostomidae) from Peru, with Taxonomic and Biogeographic Comments on Species of the Anoura caudifer Complex. Acta Chiropterologica 20:31–50.

Rivas-Pava, M., H. Ramírez-Chaves, Z. álvarez, and B. Niño-valencia. 2007. Catálogo de los mamíferos presentes en las colecciones de referencia y exhibición del Museo de Historia Natural de la Universidad del Cauca. Taller Editorial Universidad del Cauca, Popayán.

Schneider, C. A., W. S. Rasband, and K. W. Eliceiri. 2012. NIH Image to ImageJ: 25 years of image analysis. Nature methods 9:671.

Solari, S., Y. Muñoz-Saba, J. V. Rodríguez-Mahecha, T. R. Defler, H. E. Ramírez-Chaves, and F. Trujillo. 2013. Riqueza, endemismo y conservación de los mamíferos de Colombia. Mastozoología neotropical 20:301–365.

Solari, S., V. Pacheco, and E. Vivar. 1999. Nuevos registros distribucionales de murciélagos peruanos. Revista Peruana de Biología 6:152–159.

Soriano, P. J., A. Ruiz, and A. Arends. 2002. Physiological Responses to Ambient Temperature Manipulation by three Species of Bats from Andean Cloud Forests. Journal of Mammalogy 83:445–457.

Velazco, P. M.2005. Morphological phylogeny of the bat genus Platyrrhinus Saussure, 1860 (Chiroptera: Phyllostomidae) with the description of four new species. Fieldiana Zoology:1–53.

Velazco, P. M.and B. D. Patterson. 2008. Phylogenetics and biogeography of the broad-nosed bats, genus Platyrrhinus (Chiroptera: Phyllostomidae). Molecular Phylogenetics and Evolution 49:749–759.

Velazco, P. M.and N. B. Simmons. 2011. Systematics and taxonomy of great striped-faced bats of the genus Vampyrodes Thomas, 1900 (Chiroptera: Phyllostomidae). American Museum Novitates 3710:1–35.

Wilson, D. E.and D. Reeder. 1993. Mammal species of the world. Smithsonian Institution Press, WashingtonDC.

